# Revisiting the functional properties of NPF6.3/NRT1.1/CHL1 in xenopus oocytes

**DOI:** 10.1101/244467

**Authors:** Mélanie Noguero, Sophie Léran, Eléonore Bouguyon, Chantal Brachet, Pascal Tillard, Philippe Nacry, Alain Gojon, Gabriel Krouk, Benoît Lacombe

## Abstract

Within the Arabidopsis NPF proteins, most of the characterized nitrate transporters are low-affinity transporters, whereas the functional characterization of NPF6.3/NRT1.1 has revealed interesting transport properties: the transport of nitrate and auxin, the eletrogenicity of the nitrate transport and a dual-affinity transport behavior for nitrate depending on external nitrate concentration. However, some of these properties remained controversial and were challenged here. We functionally express WT NPF6.3/NRT1.1 and some of its mutant in Xenopus oocytes and used a combination of uptake experiments using ^15^N-labelled nitrate and two-electrode voltage-clamp. In our experimental conditions in xenopus oocytes, in the presence or in the absence of external chloride, NPF6.3/NRT1.1 behaves as a non-electrogenic and pure low-affinity transporter. Moreover, further functional characterization of a NPF6.3/NRT1.1 point mutant, P492L, allowed us to hypothesize that NPF6.3/NRT1.1 is regulated by internal nitrate concentration and that the internal perception site involves the P492 residue.

---

Most nitrogen enters the food chain by plants. As a consequence, understanding nitrogen uptake and assimilation in plants is of major importance to increase nitrogen use efficiency (NUE). Nitrate is one of the most important nitrogen sources for plants in both natural and agricultural systems. Uptake of nitrate by plant roots has been studied in different species and the following scheme is commonly accepted; there are inducible or constitutive high affinity transport systems (iHATS and cHATS) and low affinity transport systems (iLATS and cLATS) responsible for nitrate transport in low (<0.5 mM) and high external nitrate concentrations (>0.5 mM), respectively. Most of the HATS is supported by members of the NRT2 transporter family (Krapp *et al.,* 2014): NRT2.1, 2.2, 2.4 and 2.5 (Noguero and Lacombe, 2016); whereas LATS is supported by members of the NPF transporter family (Léran *et al.,* 2014; Corratgé-Faillie and Lacombe, 2017). In *Arabidopsis thaliana,* the founding member of the NPF family, known as NPF6.3/NRT1.1/CHL1 (Tsay *et al.,* 1993), is an exception because it has been demonstrated to behave as a dual affinity transporter (Liu *et al.,* 1999)(Ho and Frommer, 2014). Later, this property has been demonstrated for NPF6 subfamily member in other species: in *Medicago truncatula* for MtNRT1.3/MtNPF6.8 (Morere-Le Paven *et al.,* 2011) and in maize for ZmNPF6.6 (Wen *et al.,* 2017). NPF6.3/NRT1.1/CHL1 switches between low- and high-affinity depending on the phosphorylation state of the threonine 101 (T101) that is monitored by the CIPK23 kinase and its regulating partner CBL9 (Liu and Tsay, 2003; Ho *et al.,* 2009): the HATS mode is mediated by NPF6.3 transporters phosphorylated on T101. Structural studies demonstrate that oligomerization state of the transporters is the basis of the dual affinity (Parker and Newstead, 2014; Sun *et al.,* 2014; Sun and Zheng, 2015). In addition to nitrate influx, NPF6.3 can also mediate nitrate-efflux from cells (Leran *et al.,* 2013). Moreover, the role of NPF6.3 is not limited to nitrate transport, this protein also plays an important role in nitrate sensing (Bouguyon *et al.,* 2012, 2015), by mediating auxin transport (Munos *et al.,* 2004; Remans *et al.,* 2006; Ho *et al.,* 2009; Krouk *et al.,* 2010; Gojon *et al.,* 2011; Mounier *et al.,* 2013). As well as nitrate transport, this nitrate-sensing role is regulated by phosphorylation/dephosphorylation events (Ho *et al.,* 2009; Leran *et al.,* 2015*a*).

*In planta* experiments performed on different NPF6.3 Knock-Out (KO) mutant backgrounds lead to conflicting results. Even though a dual affinity was reported early (Wang *et al.,* 1998, page 1; Liu *et al.,* 1999), only its low affinity was later confirmed (Munos *et al.,* 2004; Krouk *et al.,* 2006; Glass and Kotur, 2013). Within the framework of our characterization of NPF6.3 in xenopus oocytes, we have obtained data supporting only the NPF6.3 low affinity mode. We have also shown that, in our hands, NPF6.3 is not electrogenic and that T101 is not the only CIPK23-phosphorylated residue. Finally, combination of efflux and influx experiments using the P492L mutant transporters in xenopus oocytes indicates a regulation of NPF6.3 by intracellular nitrate concentration.

## RESULTS & DISCUSSION

### NPF6.3/NRT1.1, a non electrogenic nitrate transporter

In order to assess NPF6.3 properties in heterologous system, we expressed NPF6.3 wild-type using injection of *in vitro* transcribed cRNA into xenopus oocytes. Two-electrode voltage-clamp was used to study the transport capacities of NPF6.3 (Wagner *et al.,* 2000). This technique allows recording activity of transporters mediating net currents in response to membrane voltage changes. Such transporters are named electrogenic transporters because the transport cycle stoichiometry results in an unbalanced charge moving across the membrane thus creating an apparent current. Negative currents are mediated by net influx of cations (or net efflux of anions), whereas positive currents are mediated by net influx of anions (or net efflux of cations). Control and NPF6.3-expressing oocytes were challenged with the same voltage protocol in the absence or presence of 10 mM nitrate (Fig. 1A). Currents recorded in both oocyte batches are identical (Fig. 1B and C). These currents are mediated by endogenous oocyte transporters (Weber, 1999*a*,*b*). The histogram of currents recorded at −125mV shows an increase in the negative current in response to nitrate addition in the external media. However, this current induction by nitrate is observed in both control and NPF6.3-expressing oocytes, demonstrating that this current is not NPF6.3-dependent but mediated by endogenous transporters. Moreover, no nitrate- or NPF6.3-dependent current is recorded for +25mV. Another set of experiments was performed after substituting salts (NaCl and KCl) to mannitol in the bathing solutions. In these conditions, and within a large range of membrane potentials (from −125 mV to +25 mV), in NPF6.3-expressing oocytes, nitrate is not able to induce any currents different from the ones recorded in nitrate-free solution (Fig. 1D). Before stating about NPF6.3 transport properties, the presence of functional transporters has been assessed using nitrate labeled with the non-radioactive stable nitrogen isotope (^15^NO_3_^-^).

**Fig. 1:**
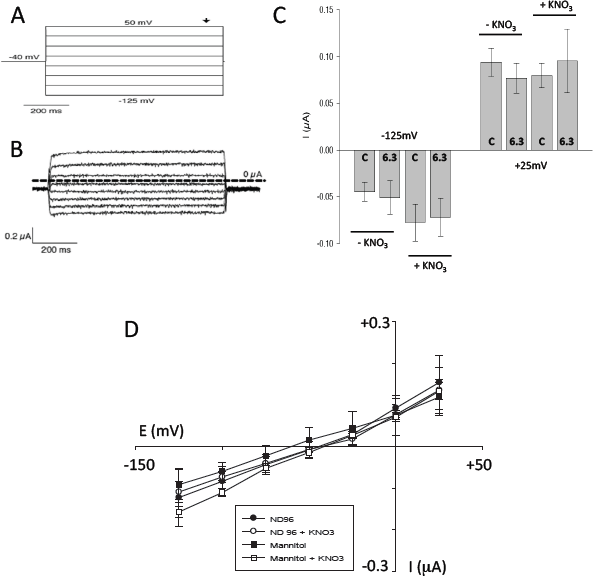
Electrophysiological characterization of NPF6.3 protein in xenopus oocytes. (A) Two-Electrode Voltage-Clamp protocol. The holding potential was −40 mV, voltage steps were applied to potentials ranging between +50 and −125 mV with 25 mV decrements. (B) Representative macroscopic current traces recorded with the two-electrode voltage-clamp technique in a 0 mM NO_3_^-^ external solution from NPF6.3-expressing oocytes. (C) Histogram of currents recorded at −125 mV and +25 mV in the absence or presence of 10 mM NO_3_^-^ in the external media, for control and NPF6.3-expressing oocytes. Data are mean +/− SE (n=10-12 oocytes). (D) I/V plots from data recorded on NPF6.3-injected oocytes. Four solutions were tested (all at pH6.0) without nitrate (black symbols) or with 10 mM nitrate (open symbols). The background solution is ND96 (circles) or manitol (squares).

### NPF6.3/NRT1.1, a low affinity nitrate transporter

We then quantified the nitrate transport activity of NPF6.3 in xenopus oocytes. We first looked at the effect of the external solution’s acidification on nitrate accumulation (Fig. 2A). Lowering pH strongly increased the accumulation demonstrating a positive effect of H+ on nitrate uptake. Since several anion transporters are able to transport nitrate as well as chloride, as ZmNPF6.4 and 6.6 (Wen *et al.,* 2017), we then remove the chloride from the external solutions and replace it by gluconate. Nitrate accumulation in both conditions is very similar suggesting that chloride is not transported by NPF6.3 (Fig.2B).

**Fig. 2:**
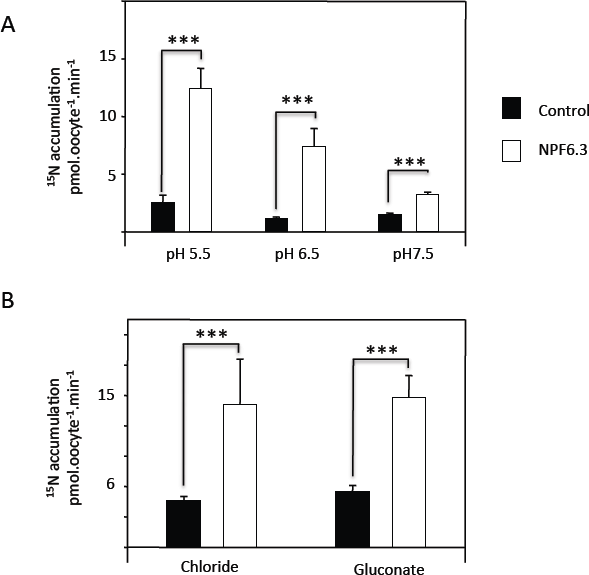
Effect of external pH an anions on ^15^N accumulation in NPF6.3-expressing oocytes. (A) ^15^N accumulation in control and NPF6.3-expressing oocytes at different external pH (5.5, 6.5 and 7.5). Data are mean +/− SE (n=7 oocytes, batched by 2), ***P<0.001, two-sided t-test after comparison with control oocytes. (B) ^15^N accumulation in control and NPF6.3-expressing oocytes with different external anions. The bathing solution is ND96 with chloride or gluconate. Data are mean +/− SE (n=6 batches of 2 oocytes), ***P<0.001, two-sided t-test after comparison with control oocytes.

Then, the effect of a large range (0 – 30 mM) of external nitrate concentrations on ^15^N accumulation in oocytes was tested (Fig. 3A and B). Within this range of concentrations, a Michaelis-Menten fit leads to a Km of 9 mM, a value close to the previously published ones (8 mM (Huang *et al.,* 1996); 4 mM (Liu *et al.,* 1999)) and demonstrates a low-affinity nitrate transport capacity. However, a closer look at the 0-1 mM concentrations range (Fig. 3b) does not support the dual-affinity model previously reported (Wang *et al.,* 1998; Liu *et al.,* 1999; Liu and Tsay, 2003). Nitrate accumulation measurements for low external concentrations do not reach a plateau and do not fit a Michaelis-Menten equation. To avoid any confusion between HATS and LATS, we then only focus on the low affinity transport system using high external nitrate concentrations.

**Fig. 3:**
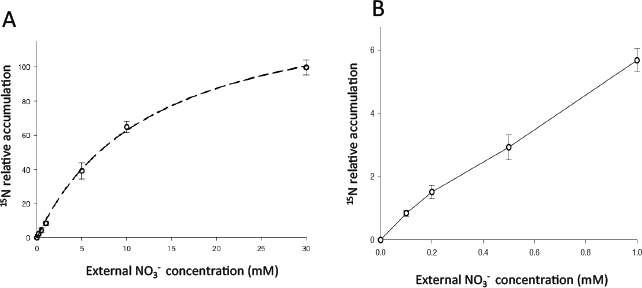
Effect of external nitrate concentration on ^15^N accumulation in NPF6.3-expressing oocytes. (A) NPF6.3-dependent ^15^N accumulation at different external nitrate concentrations range (0 – 30 mM ^15^NO_3_^-^). Data are mean +/− SE (n=10-12 oocytes, batched by 2). (B) Enlarged view of the NPF6.3-dependent ^15^N accumulation for 0 – 1 mM nitrate concentration range. Data are mean +/− SE (n=10-12 batches of 2 oocytes).

Previous work identified phosphorylated threonine on NPF6.3 (Liu and Tsay, 2003). Site directed mutagenesis and heterologous expression in xenopus oocytes identify T101 as the residue involved in switching between high- to low-affinity transport capacity of NPF6.3 (Liu and Tsay, 2003) and also as the residue being targeted by CIPK23 (Ho *et al.,* 2009). To study this regulation, we expressed the NPF6.3 native protein and two mutated forms of T101 in xenopus oocytes. The T101A substitution mimics a non-phosphorylated and non-phosphorylable form, described to be impaired in high-affinity nitrate transport, and T101D, a phosphorylated-like and non-dephosphorylable-mimic form, affected in the low-affinity nitrate transport (Liu and Tsay, 2003; Ho *et al.,* 2009). As shown by the ^15^N accumulation measurement after 2 hours incubation in 10 mM ^15^N-nitrate, both T101 NPF6.3-mutated transporters are able to transport nitrate in the low-affinity range (10 mM, Fig. 4). The difference in absolute accumulation level (e.g. T101D accumulation is higher than WT) does not give insight into the transport activity of the transporter because we have not access to the number of active transporters. Since the CIPK23 kinase - together with its regulating partner, the calcium sensor CBL9 – was identified as the kinase responsible for T101 phosphorylation (Liu and Tsay, 2003; Ho *et al.,* 2009), we tested the effect of CIPK23-CBL9 and CIPK23-CBL1 (CBL1 being also able to regulate NPF6.3/NRT1.1 in a CIPK23-dependent manner (Leran *et al.,* 2015*a*)) on nitrate transport activity (Fig. 4). It should be noticed that the cytoplasmic calcium concentration of oocyte is in a 1-10 mmolar range (Weber, 1999*a*,*b*), a concentration sufficient to activate CBLs. Both T101A and T101D mutants are still sensitive to the kinase/calcium sensor complexes. The co-expression of the native or mutated forms of NPF6.3 with CIPK23/CBL1 and CIPK23/CBL9 demonstrates that T101 is not the (only) residue involved in NPF6.3 sensitivity to CIPK23 and that the phosphorylation of these other putative residues affects NPF6.3 low-affinity transport capacity.

**Fig. 4:**
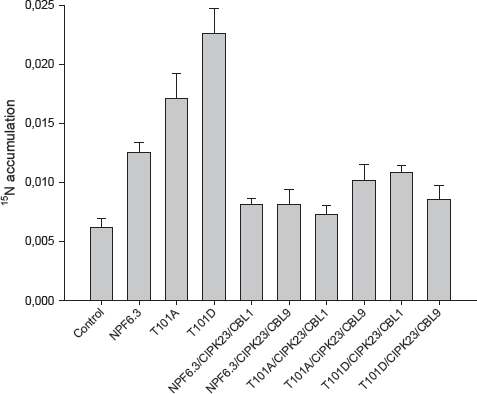
Nitrate transport activity of the native NPF6.3 and two T101 mutated forms of NPF6.3 proteins and their regulation by CIPK23-CLB1/9. ^15^N accumulation after 2 hours incubation in 10 mM ^15^NO_3_^-^, in control oocytes, native, T101A or T101D NPF6.3-expressing oocytes, with or without CIPK23/CBL1 or CIPK23/CBL9. Data are mean +/− SE (n=10-12 batches of 2 oocytes).

### NPF6.3/NRT1.1, a nitrate transporter regulated by internal nitrate

A NPF6.3 mutant defective for nitrate influx, P492L, has been characterized in xenopus oocytes (Ho *et al.*, 2009). Interestingly, plants expressing this mutated protein are impaired in root nitrate influx but can still sense nitrate. This phenotype could be due to a default in subcellular localization, since P492L-NPF6.3 is localized in internal membranes (Bouguyon *et al.,* 2015). Our data confirm that the P492L mutated form of NPF6.3, expressed in oocytes, does not mediate nitrate transport. Indeed, when tested in influx conditions (^15^N-labeled external nitrate), ^15^N accumulation in NPF6.3-P492L is as low as in control oocytes (Fig. 5a). We also assessed another functional property of the NPF6.3 transporter in xenopus oocytes, namely the nitrate efflux (Leran *et al.,* 2013). Contrary to what was expected, the P492L mutated form conserves its nitrate efflux capacity when ^15^N nitrate is injected into the oocytes: the ^15^N content after 1 hour in P492L-expressing oocytes is similar to the ^15^N content recorded in oocytes expressing the native form of the protein (Fig. 5b). This result suggests that internal nitrate gates P492L mutant. To test this hypothesis, the ^15^N accumulation measurement was repeated using influx conditions (2 hours incubation in 10 mM ^15^N-nitrate media) but after a non-labeled (^14^N) nitrate injection into oocytes. The ^15^N accumulation level in the P492L-NPF6.3 expressing oocytes is then similar to the one measured in the oocytes expressing the native NPF6.3 (Fig. 5c).

**Fig. 5.**
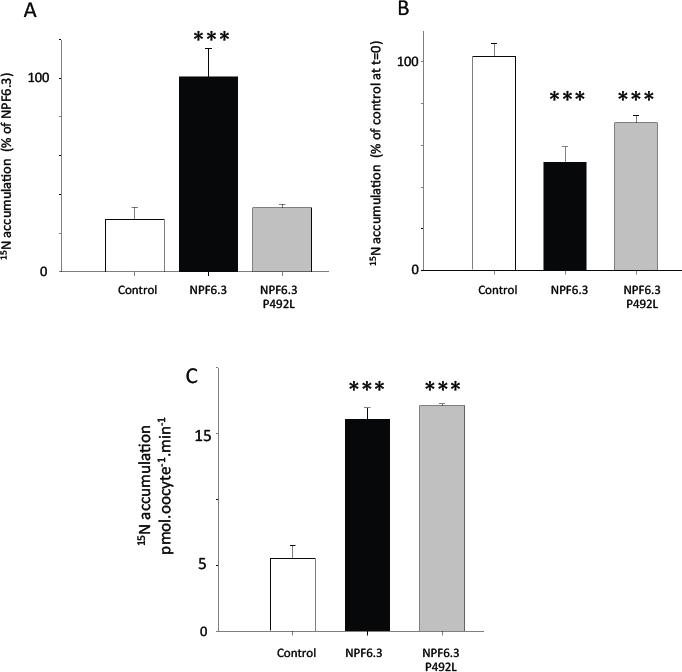
Nitrate transport activity of native NPF6.3 and P492L-NPF6.3 and their regulation by internal nitrate. (A) ^15^N accumulation in control, native NPF6.3 and P492L-NPF6.3-expressing oocytes after 2 hours incubation in a 10 mM ^15^NO_3_^-^ solution, expressed in percentage of ^15^N accumulation in NPF6.3-expressing oocytes. Data are mean +/− SE (n=10-12 batches of 2 oocytes), ***P<0.001, two-sided t-test after comparison with control oocytes. (B) ^15^N content in control, NPF6.3 and NPF6.3-P492L expressing oocytes, 1 hour after ^15^NO3^-^ injection in oocytes. Values are expressed in percentage of ^15^N remaining in control oocytes. Data are mean +/− SE (n=10-12 batches of 2 oocytes), ***P<0.001, two-sided t-test after comparison with control oocytes. (C) ^15^N accumulation in control, NPF6.3 and NPF6.3-P492L-expressing oocytes, after injection of non-labeled (^14^N) nitrate and 2 hours incubation in 10 mM ^15^NO_3_^-^. Data are mean +/− SE (n=10-12 batches of 2 oocytes), ***P<0.001, two-sided t-test after comparison with control oocytes.

Our results show that P492L-NPF6.3 activity is gated by intracellular nitrate concentration, a property that, combined with its modified sub-cellular localization, is expected to be involved in its nitrate-sensing role. In the time frame of our influx experiments (2 hours), the oocyte intracellular concentration increases from 0 to <1 mM, due to the presence of endogenous transporter, able to import nitrate (Weber, 1999*a*,*b*). This low nitrate concentration is enough to activate the NPF6.3-P492L mutated protein but not the native one.

## CONCLUSION

Despite their acknowledged usefulness, heterologous systems are known to have some drawbacks and differences in transporter’s functional properties from batch to batch and lab to lab, as already reported (Dreyer *et al.,* 1999). Even if our experiments do not allow to definitively stating about the transport properties of NPF6.3, they support a model of an electroneutral symport protein, co-transporting one nitrate ion with one proton. Furthermore, our data obtained using xenopus oocyte as a heterologous system profusely support the NPF6.3 low-affinity transport activity. However, when testing a wide range of low external nitrate concentrations we could not confirm the NPF6.3 high-affinity transport capacity. Moreover in our conditions, the NPF6.3-T101 mutants, previously used to prove the dual affinity transport system, do not present the expected nitrate transport activity regarding their phosphorylation state.

Our work on the mutated P492L form of NPF6.3 in xenopus oocytes allowed us to propose that NPF6.3 nitrate transport is gated by intracellular nitrate. This regulation should now be assessed *in planta* to figure out its physiological role.

## EXPERIMENTAL PROCEDURES

### Expression in xenopus oocytes

NPF6.3 (WT and mutants), CIPK23, CBL1 and CBL9-pGEM were linearized and *in vitro* transcribed with mMessage mMachine T7 Ultra Kit following manufacturer protocol (Life Technologies). Oocytes were obtained and injected as previously described (Lacombe and Thibaud, 1998).

### Electrophysiological recordings

Electrophysiological measurements were made 3 days after cRNA injection in oocytes, using a two-electrode voltage-clamp technique. The following bathing solutions with or without 10 mM KNO_3_ were used: ND96 (2 mM KCl, 96 mM NaCl, 1 mM MgCl_2_, 1.8 mM CaCl_2_, 5 mM MES/Tris, pH 6.0) or Mannitol (220 mM Mannitol, 1 mM MgCl_2_, 1.8 mM CaCl_2_, 5 mM MES/Tris, pH 6.0). Current passing and voltage-recording electrodes were filled with 3 M KCI and had, respectively, 0.5-1 MW and 1.5-2 MW tip resistance in 100 mM KCI. The voltage-clamp amplifier was an Axoclamp-2A (Axon Instruments, USA). Voltage-pulse protocols, data acquisition and data analysis were performed using the pClamp program suite (Axon Instruments, USA). Both membrane potential and current were recorded. Correction was made for voltage drop through the series resistance of the bath and the reference electrode by using a voltage-recording microelectrode in the bath close to the oocyte (the potential of the bath electrode was subtracted from the one of intracellular electrodes in the amplifier, allowing a real-time series resistance correction). All experiments were performed at room temperature (20-22°C).

### ^15^N accumulation measurements

Briefly, 3 days after cRNA injection, injected and control (non-injected) oocytes were incubated 2 hours in 2 ml of ND96 medium (pH 6.5) containing between 0.1 and 30 mM of ^15^N-nitrate (atom % ^15^N abundance: 99,9%) to test the influx activity (Leran *et al.,* 2015*b*). For pH experiments, ND96 solution was adjusted at pH 5.5 (MES), 6.5 (MES) or 7.5 (HEPES). For anion experiments, chloride was replaced by gluconate. Oocytes were then washed 5 times in 15 ml of ND96 medium (pH 6.5) at 4°C. For the efflux conditions, ^15^N-nitrate was injected into the oocytes, and they were washed after 1 hour incubation in a nitrate-free media as previously described (Leran *et al.,* 2015*b*).

Batches of 2 oocytes were then analyzed for total N content and atom % ^15^N abundance by Continuous-Flow Mass Spectrometry, using an Euro-EA Eurovector elemental analyzer coupled with an IsoPrime mass spectrometer (GV instruments, Crewe, UK).

## Acknowledgments

This work was supported by the Institut National de la Recherche Agronomique (CJS PhD Fellowship to SL & Projet Département BAP, BAP2013-33-NITSE to BL), Agence Nationale de la Recherche (ANR-11-JSV6-002-01-NUTSE and ANR-14-CE34-0007-01-HONIT to BL) and the Région Languedoc-Roussillon (Chercheur d’Avenir to BL).

## List of author contributions

MN, SL, EB, CB, PT, GK, BL performed research; MN, SL, EB, AG, GK, BL analyzed data; BL designed the research; MN, SL & BL wrote the paper with contributions of all authors.

